# Multi-environmental population phenotyping suggests the higher risk of wheat *Rht-B1b* and *Rht-D1b* cultivars in global warming scenarios

**DOI:** 10.1101/2022.07.18.500398

**Authors:** Zihui Liu, Zunkai Hu, Zhiwei Zhu, Junmei Cao, Jialiang Zhang, Xiangyu Ma, Xinzhong Zhang, Xiaoming Wang, Wanquan Ji, Shengbao Xu

## Abstract

After six decades practice, the semi-dwarf alleles *Rht-B1b* and *Rht-D1b* (also called *Rht-1* and *Rht-2*) had been applied into around 70% current wheat cultivars, laid the foundation for the worldwide wheat production supply, then the agronomic traits controlled by the two alleles still keep unclear except dwarfing wheat. Here 13 agronomic traits were investigated in 400 wheat accessions with seven environments, uncovered the genetic effects of *Rht-B1b* and *Rht-D1b* on wheat structure and yield traits in different genetic backgrounds and environments, and the distinct genetic effects between *Rht-B1b* and *Rht-D1b*, suggesting that the introduction of green revolution alleles profoundly shaped the agronomy traits of modern wheat cultivars. The late-sowing assays and regression analysis based on the phenotypic and their meteorological data showed the accessions containing *Rht-B1b* and *Rht-D1b* are more sensitive to the temperature increase, and the *Rht-D1b* may lost additional 8% yield compared the cultivars without the green revolution alleles if the average temperature increases 1 °C. These results suggest the application of green revolution are facing more challenges to maintain the futural wheat production supply in global warming scenarios.

---

The introduction of the semi-dwarfing trait into wheat cultivars during the 1960s and 1970s, confers great lodging resistance, allowing high fertilizer input, thereby significantly enhance yield of crop, was a defining as the “Green Revolution” ^1,2^. The Green Revolution alleles in wheat include the *Rht-B1b* and *Rht-D1b* (also named as *Rht-1* and *Rht-2*), which introduce premature stop codons in the N-terminal coding region of DELLA proteins to alter the GA signaling ^3,4^. Approximately 70% of cultivars in a worldwide winter wheat panel that were released in the 21st century carry one of these two “Green Revolution” alleles ^5^.

After sixty years application, people found the green revolution alleles not only function in reducing the plant height, but also involved in a broad of agronomic traits ^6,7^. However, limited by few genetic backgrounds and environmental conditions, the agronomic traits controlled by green revolution alleles showed inconsistent conclusions in previous studies ^8-10^. In addition, two green alleles, the *Rht-B1b* and *Rht-D1b*, showed minor difference in controlling wheat traits ^8,11,12^, which may related their adoption to the distinct wheat region in USA ^13^, Europe ^14,15^ and China ^16,17^, then the underlying difference still keeps unknown. These studies suggest the agronomic phenotypes controlled by *Rht-B1b* and *Rht-D1b* are far from fully dissected, which obstacles the study on underlying regulatory mechanism and its precisely application in wheat breeding and production.

In addition, current practices had demonstrated that the wheat cultivars containing *Rht-B1b* and *Rht-D1b* are more vulnerable to drought stress ^18,19^, indicating a potential weakness of *Rht-B1b* and *Rht-D1b* in adapt to environmental stress, which may influence the application of green revolution alleles in futural global warming scenario. As the most adopted wheat alleles, their adaptation to high temperature has to be evaluated to ensure wheat production security for the future.

Here, 13 agronomic traits of *Rht-B1b* and *Rht-D1b* were investigated in seven environments across three years on 400 wheat accessions, demonstrating the global genetic effects of *Rht-B1b* and *Rht-D1b* on wheat traits and its environmental stability, providing the reliable phenotypes and its occurrence probability in field for the precisely application in current and futural wheat breeding. The heat sensitivity of green revolution alleles suggests current wheat breeding and production have to adjusted to face the global warm scenarios.

## The bread wheat population for agronomic phenotyping

A total 400 wheat accessions were collected from all over the world, and were genotyped by RNA-seq ^20^ and confirmed by the KASP assay, including 105 *Rht-B1b* (*B1b)* accessions, 98 *Rht-D1b* (*D1b*) accessions, and 197 accessions without *B1b* and *D1b* (refer to *NO* thereafter) (Figure 1A). Thirteen agronomic traits were investigated in seven environments across three years (Table S1). The correlations analysis suggests the plant height in this population is significantly correlated with peduncle length, total tiller number, kernel number per spike and thousand kernel weight (Figure 1B), indicating that these traits may be affected by the introduction of green revolution alleles.

**Figure. 1.**
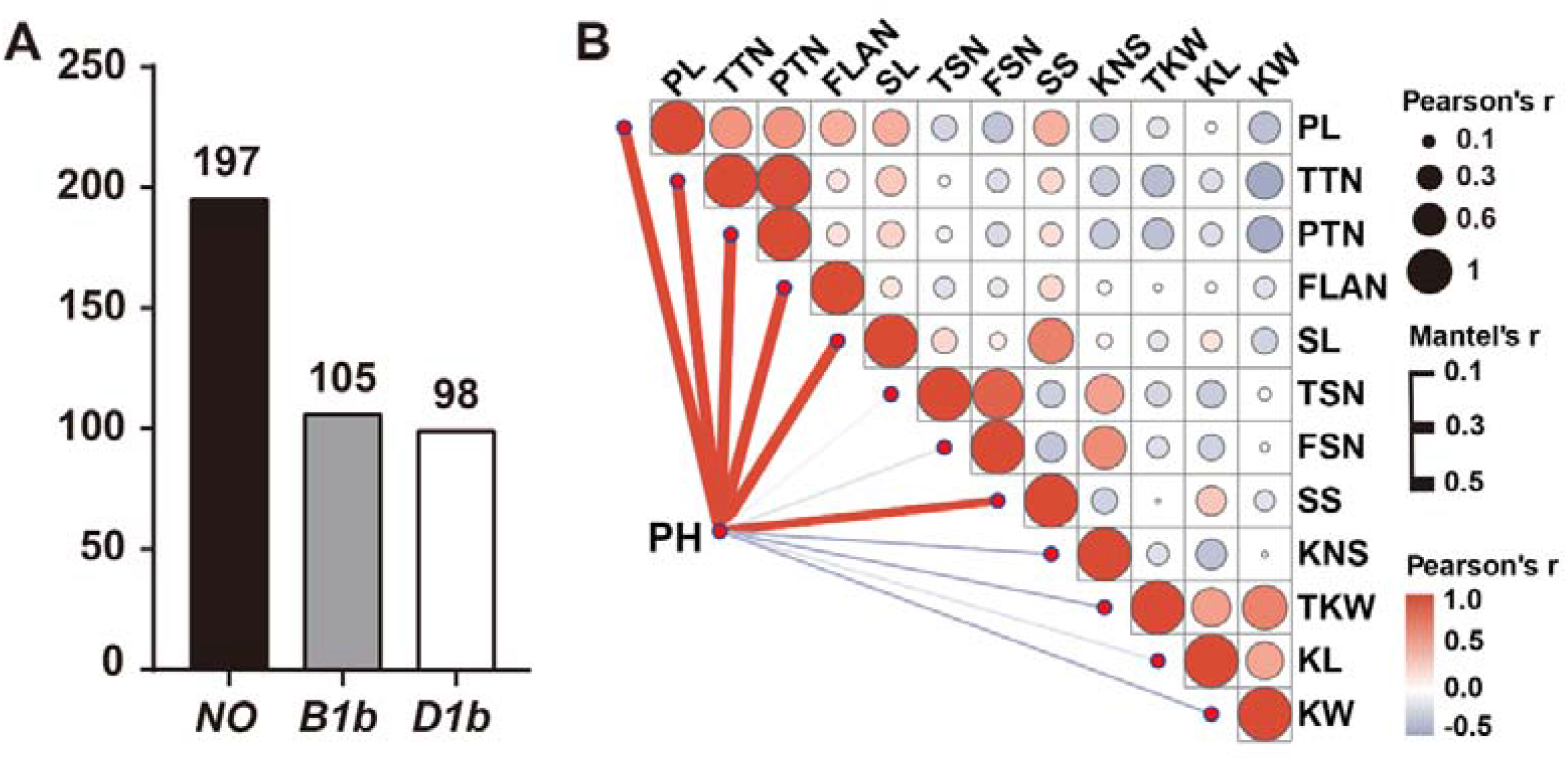
The wheat population and agronomic traits. (A) The number of wheat accessions with different genotypes in the population. *NO*, wheat accessions without *Rht-B1b* and *Rht-D1b*; *B1b*, wheat accessions containing *Rht-B1b*; *D1b*, wheat accessions containing *Rht-D1b*. The number indicates the accessions number in each genotype. (B) The correlations analysis between different agronomic traits by the best linear unbiased estimate (BLUE) values. PH, plant height; PL, peduncle length; TTN, total tiller number; PTN, productive tiller number; FLAN, flag leaf angle; SL, spike length; TSN, total spikelet number; FSN, fertile spikelet number; SS, spikelet spacing; KL, kernel length; KW, kernel width; KNS, kernel number per spike; TKW, thousand kernel weight. Red lines mean positive correlation and gray lines mean negative correlation. The correlations were analyzed by the “corrplot” in the R package using the BLUE values in Table S1.

## The genetic effects of *Rht-B1b* and *Rht-D1b* on wheat plant structure and yield

The BLUE (best linear unbiased estimator) value of agronomic traits in seven environments was used to evaluate the difference between each genotype, and 1000 permutation tests were performed on the 50% random sampling combination in each group to evaluate genomic background independence, termed as genomic probability (GP). We found that all the plant structure related traits are significantly different between *NO* and dwarf group (the *B1b* and *D1b*), and also significantly different between *B1b* and *D1b* with high GP (> 0.75 in Figure 2). Notably, the *Rht-D1b* generally displayed a stronger effect in altering structure traits, consistent with the previous observation that the dwarf effect from *D1b* is significantly and stably higher than that in *B1b* ^5^.

**Figure. 2.**
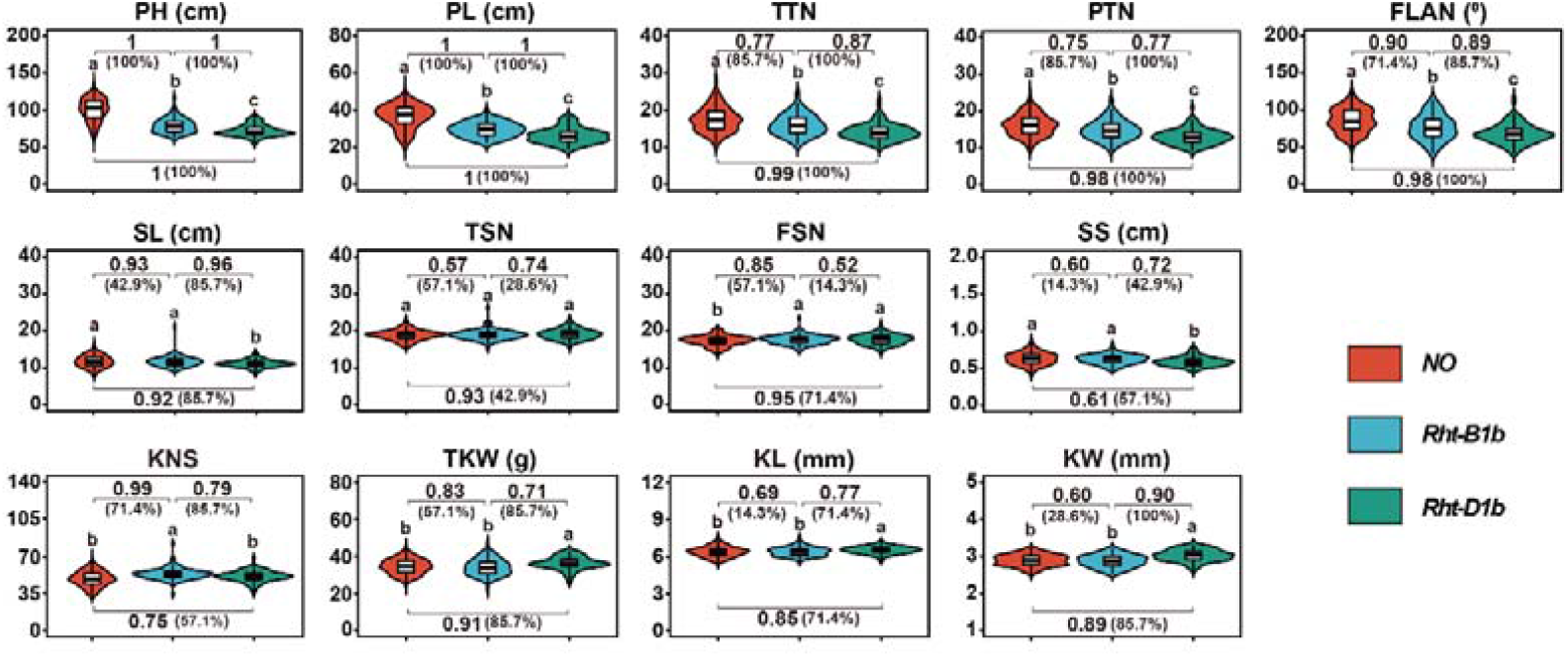
The function of the *Rht-B1b* and *Rht-D1b* on wheat traits. The *Rht-B1b* and *Rht-D1b* were compared to that of *NO* accessions, the letters at the top of violin plots indicate the significant difference between each group (two-sided Dunn’s Kruskal– Wallis test, *P* < 0.01). The permutation tests (1000) were performed on the 50% random samples combination in each group, the numbers represent the ratio of comparison showing significantly difference / all comparison (1000). The percentages in parentheses indicate environmental probability, the number of environments that have significant difference effect / the total number of environments (7). The data come from the BLUE values (n=400) of seven environments, as shown in Table S1.

However, the general significant difference can’t be detected on the spike and kernel related traits between the dwarf and *NO* group (Figure 2). Interestingly, the *D1b* confers higher kernel length, kernel width and final kernel weight, result in 2.1 microgram increase for each kernel (Figure 2), equal to 6% yield increase compared to the *NO*. In contrast, the *B1b* showed higher kernel number per spike, which result in 4.2 kernels increase for one spike, contribute around 8% yield increase compared to that of *NO*. The significantly difference of kernel number and kernel weight are also observed between the cultivar group of *B1b* and *D1b* with medium GP (0.5-0.8), highlighting that the distinct positive effect of *Rht-B1b* and *Rht-D1b* on wheat yield, providing new insights for higher yield conferred by green revolution alleles ^1,18^.

## Environmental variations of Green Revolution genes

To evaluate the environmental stability of the effect of green revolution alleles, the environmental probability (EP), which present the occurrence probability with significant difference in investigated environments, was created to describe the environmental stability (Figure 2, Table S1). We found the effect of green revolution alleles were stable in plant structure traits with high EP, especially *D1b*, which showed the significant difference compared to *NO* group in more than five environments out of all seven investigated environments, indicating that the effects of green alleles on wheat structure traits is relative stable in different environments.

For spike related trait, only the kernel number per spike controlled by *B1b* showed high environmental stability (>70%), while the effect of *D1b* on spike length, thousand kernel weight and kernel width showed medium environmental stability (≥ 43.9%) (Figure 2). These results suggested the roles of green revolution alleles in kernel traits are sensitive to the environments.

## Functional verification of *Rht-B1b* and *Rht-D1b*

To evaluate the pleiotropism of *B1b* and *D1b*, the GWAS with resample model inclusion probability were performed on of the wheat traits, showing six and seven traits were significantly associated with *B1b* and *D1b* respectively in this nature population (Table 1, Table S2), supporting the pleiotropism of *Rht-B1b* and *Rht-D1b* in regulating the wheat traits (Table 1). Considering different environmental response of wheat accessions is also derived from the genomic difference, we combined the EP and the GP, and we found the green alleles could be identified in 91% wheat traits in GWAS if the combined probability is over 0.7 in given trait (Table 1).

**Table 1.** Verification of *Rht-B1b* and *Rht-D1b* for all plant structure and yield related traits.

| Traits | *Rht-B1b* |  |  |  |  | *Rht-D1b* |  |  |  |  |
| --- | --- | --- | --- | --- | --- | --- | --- | --- | --- | --- |
|  | EP (%) | GP | CP (%) | PE (%) | GWASP (%) | EP (%) | GP | CP (%) | PE (%) | GWASP (%) |
| PH | 100 | 1 | 100 | -22.6* | 100 | 100 | 1 | 100 | -27.7* | 100 |
| PL | 100 | 1 | 100 | -19.7* | 85.7 | 100 | 1 | 100 | -27.9* | 85.7 |
| TTN | 85.7 | 0.77 | 66.0 | -8.65* | 0 | 100 | 0.99 | 99.0 | -19* | 14.3 |
| PTN | 85.7 | 0.75 | 64.3 | -8.09* | 0 | 100 | 0.98 | 98.0 | -18.3* | 42.9 |
| FLAN | 71.4 | 0.90 | 64.3 | -10.9* | 0 | 100 | 0.98 | 98.0 | -19.8* | 85.7 |
| SL | 42.9 | 0.93 | 39.9 | -0.42 | 0 | 85.7 | 0.92 | 78.8 | -5.1* | 0 |
| TSN | 57.1 | 0.57 | 32.5 | +0.37 | 0 | 42.9 | 0.93 | 39.9 | +0.69 | 0 |
| FSN | 57.1 | 0.85 | 48.5 | +1.9* | 0 | 71.4 | 0.95 | 67.8 | +1.53* | 0 |
| SS | 14.3 | 0.60 | 8.60 | -1.59 | 0 | 57.1 | 0.61 | 34.8 | -6.35* | 0 |
| KNS | 71.4 | 0.99 | 70.7 | +8.95* | 100 | 57.1 | 0.75 | 42.8 | +3.47 | 0 |
| TKW | 57.1 | 0.83 | 47.4 | +2.59 | 71.4 | 85.7 | 0.91 | 78.0 | +6.32* | 57.1 |
| KL | 14.3 | 0.69 | 9.90 | +0.16 | 57.1 | 71.4 | 0.85 | 60.7 | +2.2* | 0 |
| KW | 28.6 | 0.60 | 17.2 | +1.04 | 71.4 | 85.7 | 0.89 | 76.3 | +4.5* | 28.6 |

EP, Environmental probability, presents the ratio of the number of environments showing significant difference / all investigated environments (seven environments);
GP, Genetic probability, presents the ratio of comparison showing significantly difference / all comparison (the 1000 permutation);
CP, Combined probability, presents the probability of EP \* GP;
PE, Phenotype effect, presenting the effect of dwarf allele on the given traits;
GWASP, GWAS probability, presents the number of environments detecting dwarf allele / all environments in GWAS analysis.

The pleiotropism of green alleles was further supported by three F_2_ populations constructed by the accessions with *B1b* and *D1b* respectively (Figure 3), and support the function of *B1b* in reducing SL (Figure 3) although the SL displayed a low combined probability (Table 1). Moreover, F_2_ population showed that the *B1b* also has a role in enhancing TKW, but its effect is much lower than that of *D1b* (Figure 3, Table 1).

**Figure 3.**
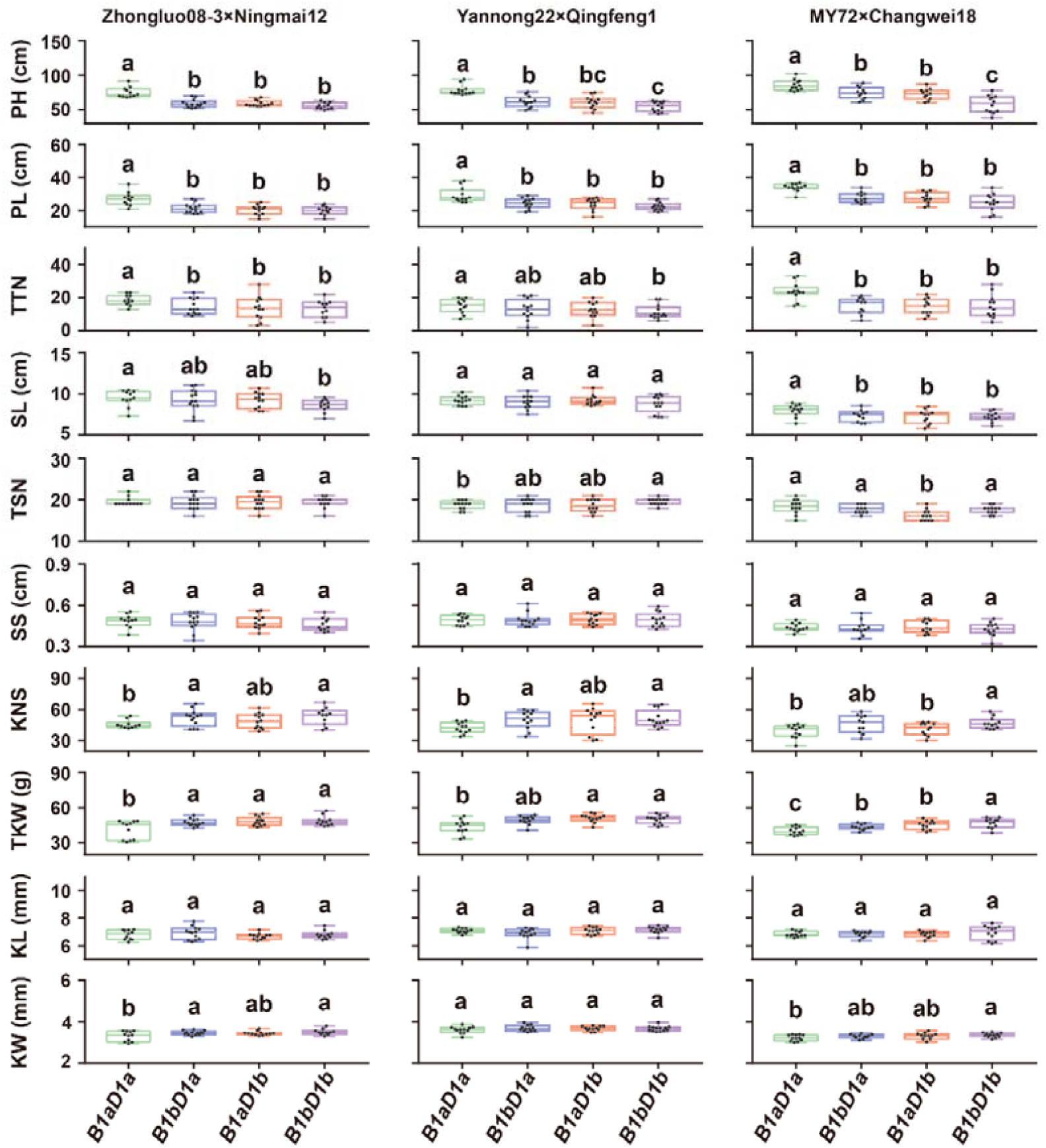
Validation on the genetic role of the *Rht-B1b* and *Rht-D1b* in three F2 populations. The parents are Zhongluo08-3 (*Rht-D1b*); Ningmai12 (*Rht-B1b*), Yannong22 (*Rht-B1b*); Qingfeng1 (*Rht-D1b*); MY72 (*Rht-B1b*) and Changwei18 (*Rht-D1b*), respectively. Different letters indicate statistical significance, determined using two-sided Dunn’s Kruskal–Wallis test (*P*□<□0.05). *B1aD1a* refers *Rht-B1a* & *Rht-D1a*; *B1bD1a* refers *Rht-B1b* & *Rht-D1a*; *B1aD1b* refers *Rht-B1a* & *Rht-D1b*; *B1bD1b* refers *Rht-B1b* & *Rht-D1b*.

These investigations suggest the introduction of *Rht-B1b* and *Rht-D1b* substantially altered multitude traits in current wheat cultivars by their genetic effect. However, the alterations conferred by green revolution alleles occur in field with different probability for the different genomic background.

## Spatiotemporal application of the Green Revolution genes in China

In our wheat population, including 87 landraces (LA) and 255 historical/modern cultivars (MC) (Table S1). We found the application frequency of *B1b* and *D1b* dramatically increase in MC (Figure 4A). The *D1b* was preferentially adopted by the Chinese cultivars in the Yellow and Huai wheat zone (50.4%), while the *Rht-B1b* was more adopted by Chinese cultivars in Yangtze River winter wheat zone and the introduced cultivars from abroad (Table S3), consistent to that there is a distinct environmental adoption of two green revolution genes ^15,21,22^. The distinct preferentially adoption of *B1b* and *D1b* displayed a clearer trend in the CMC released after 2000. The total application frequency of *B1b* and *D1b* increased from 45.9% to 78.1% in CMC, which is released before and after 2000 (Figure 4B), highlighting the green revolution alleles still are the major dwarf alleles applied in current wheat varieties.

**Figure. 4.**
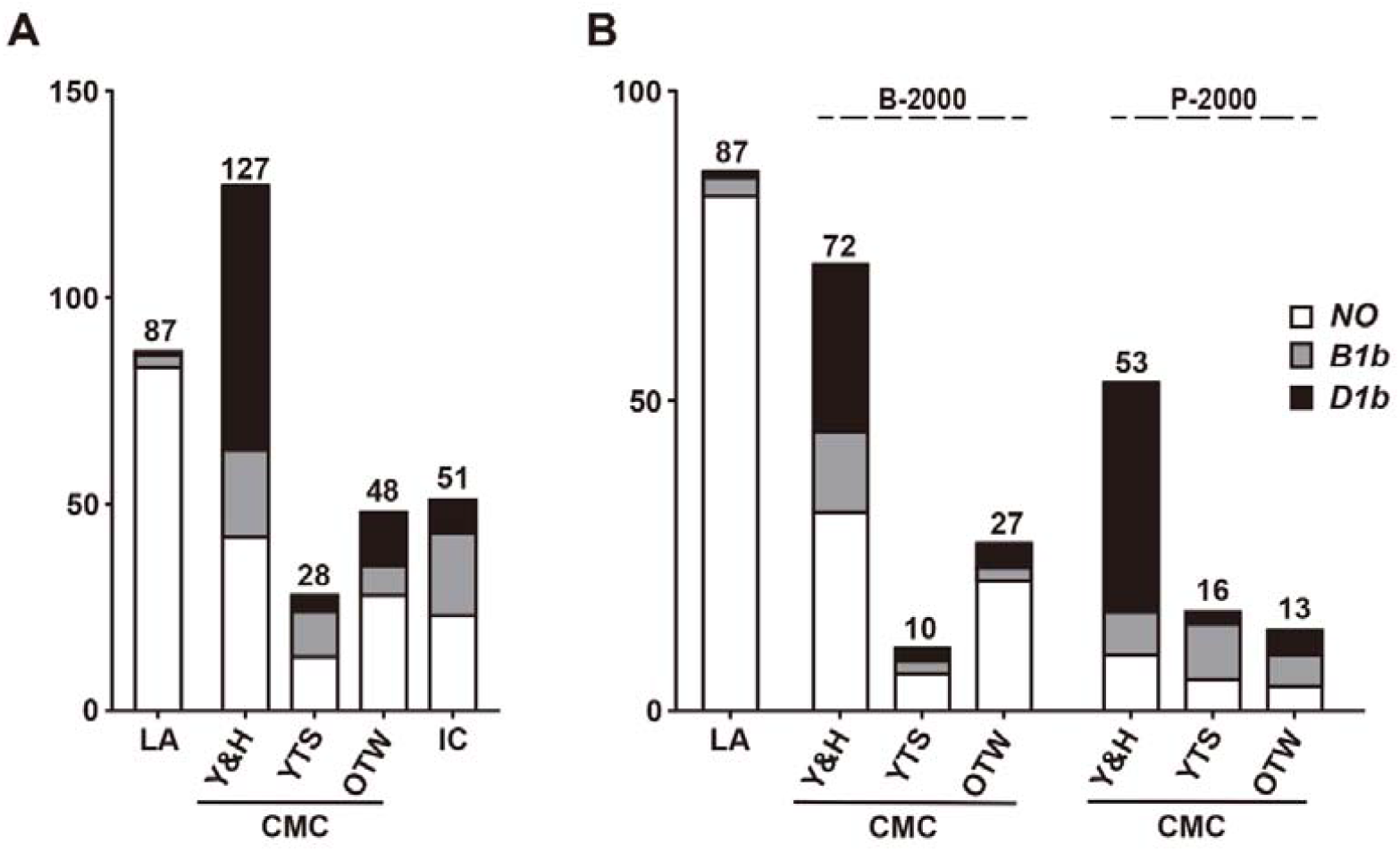
The spatiotemporal distribution of the *Rht-B1b* and *Rht-D1b*. (A) The distribution of the *Rht-B1b* and *Rht-D1b*. LA, landraces; Y&H, Yellow and Huai winter wheat zone; YTS, Yangtze River winter wheat zone; OTW, other wheat zone in China; IC, Introduced worldwide cultivars to China. The numbers in the bar indicate the number of wheat accessions, respectively. (B) The application frequency of the *Rht-B1b* and *Rht-D1b* before and after 2000. B-2000, the cultivars was released before 2000; P-2000, the cultivars was released post 2000.

## The heat adaptation potential of *Rht-B1b* and *Rht-D1b*

To evaluate the thermotolerance of green revolution alleles for the global warming challenges, late-sowing field assays ^23,24^ were performed in three years, which confers higher temperatures for different developmental stages of wheat (Figure 5A, Table S5).

**Figure. 5.**
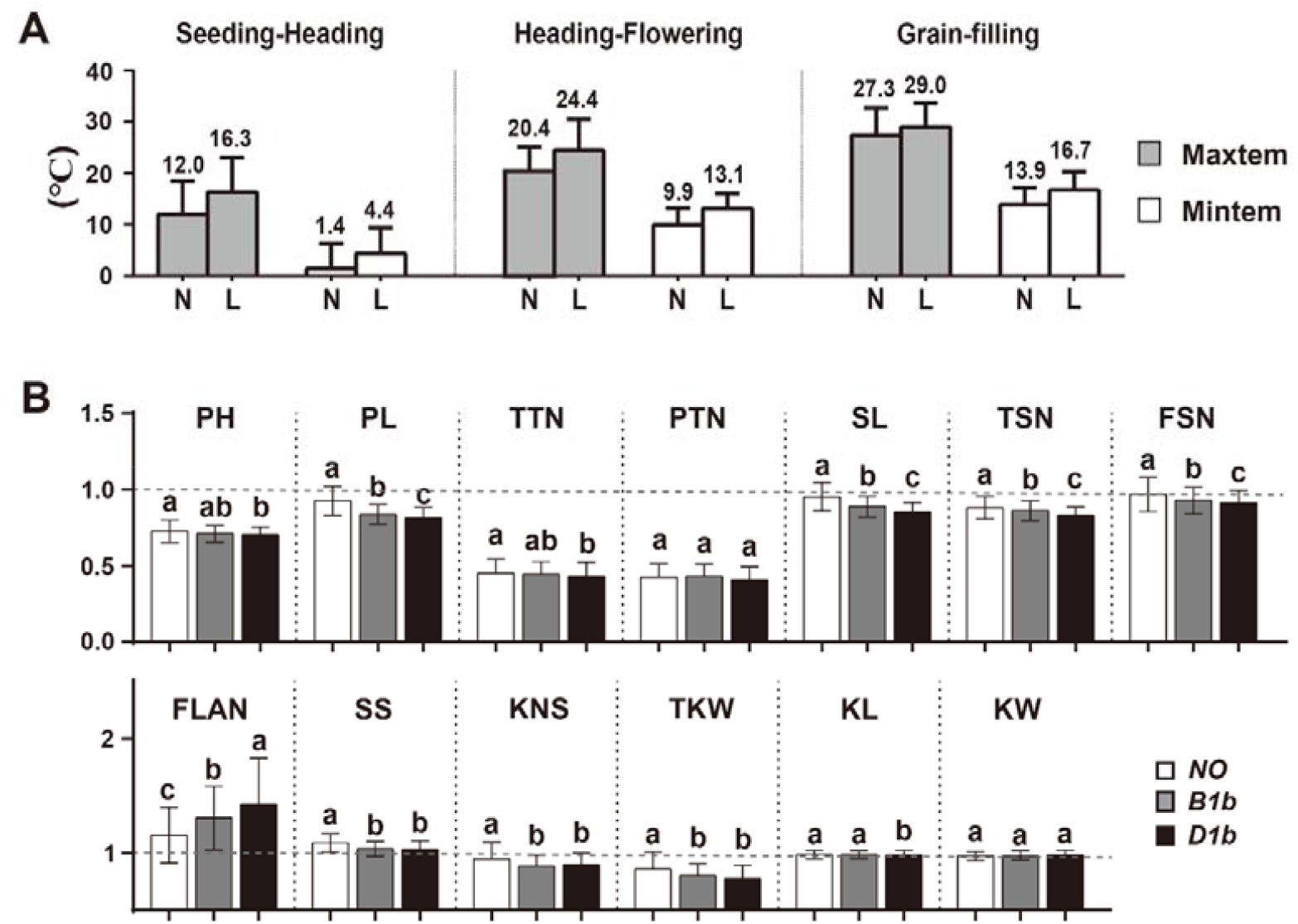
The phenotypic variations with late-sowing. (A) The temperature variations with late-sowing. N, normal-sowing; L, late-sowing. The sowing date and temperature was shown in Table S4 and S5. (B) The phenotypic alteration with late-sowing. The data is the phenotypic data in late-sowing / phenotypic data in normal-sowing. Different letters on the bars indicate the significant differences (*P* < 0.05, two-sided Dunn’s Kruskal–Wallis test), as shown in Table S6. The gray dash lines indicate the value 1.

We found that almost investigated wheat traits significantly changed by the late-sowing (Figure 5B, Table S6). For plant structure, the plant height was decreased about 30%, and the tiller number were decreased more than 50% in late-sowing condition. However, for yield traits, only the thousand kernel weight were decreased more than 15%, indicating that the vegetative growth of wheat was more sensitive to the late sowing (Figure 5B, Table S6). Notably, the agronomic traits of green revolution alleles accessions showed more severe influences by the late-sowing (Figure 5B), indicating the accessions with green revolution alleles may be more sensitive to the temperature increase than accessions in *NO* group, especially the accessions containing *Rht-D1b*.

## The effect of green alleles in response to increasing temperature

To further evaluate the effects of high temperature on *Rht-B1b* and *Rht-D1b*, the meteorological data, including the air temperature, sunlight hours, relative humidity and solar radiation in seven environments was collected (Table S7, S8), then integrate the phenotypic data from the seven environments for a multivariate regression analysis ^25,26^, and we found the temperature in different developmental stage contributes the substantial major effect to all traits of wheat population (Table S9).

Next, the average temperatures at different developmental stage were selected for the regression analysis (Table S10). With the regression equations, we predict the traits alterations when the temperature increases 1 °C in whole growth season, showing the tilling number, flag leaf angle, grain weight and plant height are the most temperature sensitive traits (Figure 6A), and the cultivars containing *B1b* and *D1b* were more sensitive to the increasing temperature compared to *NO* accessions, generally consistent to the estimation by the late sowing (Figure 5B).

**Figure. 6.**
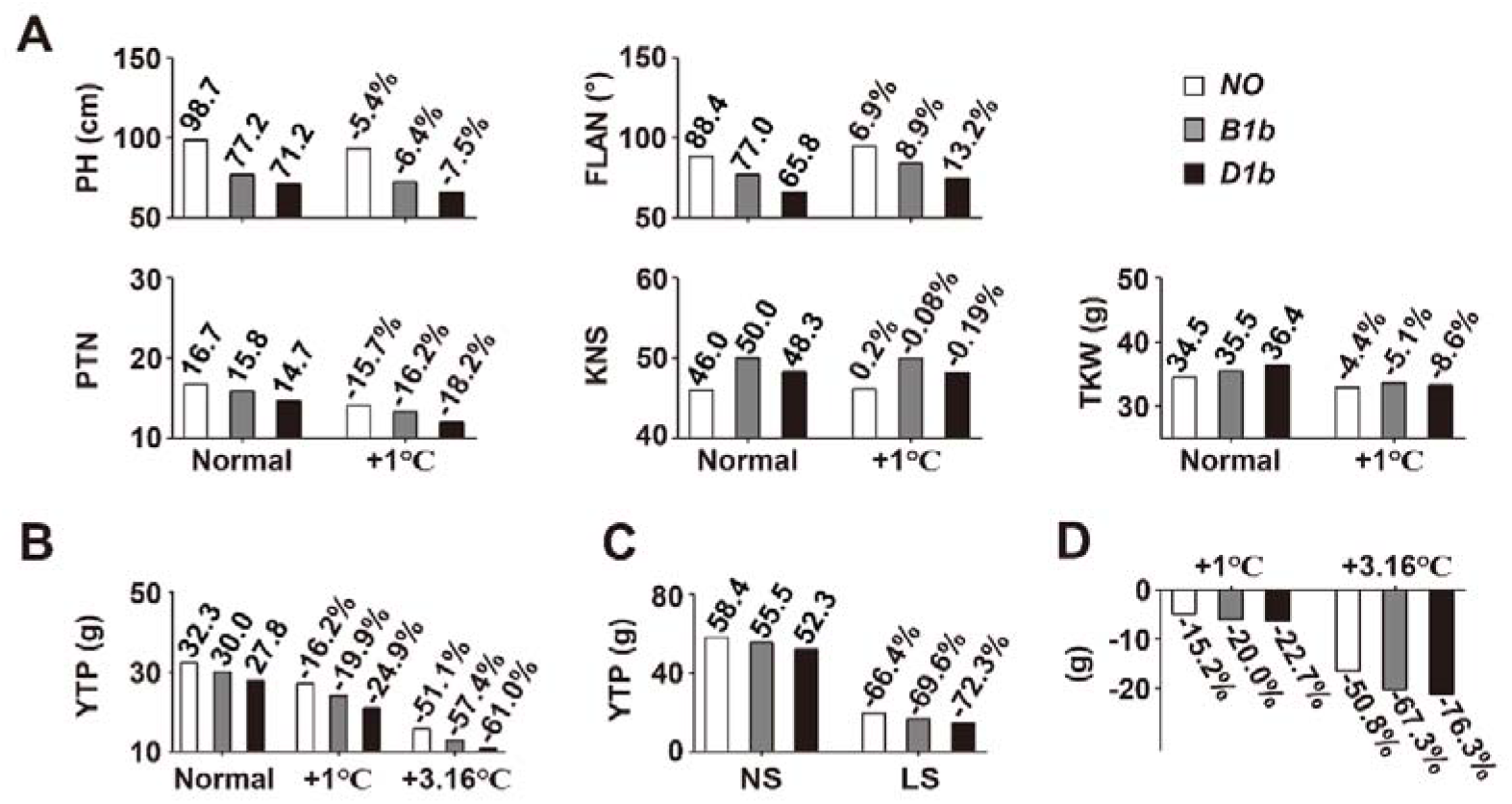
The phenotypes of the *Rht-B1b* and *Rht-D1b* in response to temperature increase. (A) The alterations of wheat traits when the temperature increased 1°C. Normal, the normal temperature of wheat growth; +1°C indicates the average temperature increase 1°C of all developmental stage. The number indicates the phenotype calculated from the regression equation with normal temperatures; the percentages indicate the phenotypic changed ratio of corresponding genotype wheat accessions when the average temperature increased 1°C. (B) The predicted yield per plant (YPP) with temperature increase by regression analysis. The YPP was estimated by productive tiller number × kernel number per spike × kernel weight. 3.16°C indicates the average temperature increase of late-sowing compared to normal-sowing. (C) The YPP with normal-sowing and late-sowing. The number indicates the YPP of normal-sowing, the percentages indicate the YPP changed ratio in late sowing compared with normal sowing. NS, normal-sowing; LS, late-sowing. (D) The variations of YPP according to the temperature variations. The percentages indicate the YPP changed ratio of corresponding genotype wheat accessions when the average temperature increased 1°C and 3.16°C.

The yield per plant (YPP) will reduce around 20% if the temperature increases 1 °C (Figure 6B). Compared with *NO* cultivars, the YPP of cultivars containing *B1b* and *D1b* showed an additional 3.7% and 8.7% penalty, respectively, if the temperature increases 1 °C (Figure 6B). Meanwhile, the late sowing in three years contribute an average of 3.16 °C temperature increase, contribute 66.4 - 72.3% penalty in YPP (Figure 6C), and an additional 3.2% and 5.9 % YPP penalty were observed in the cultivars containing *B1b* and *D1b* respectively, which showed the similar decreased trend with the predicted result (6.3% and 9.9% for *B1b* and *D1b* respectively) by the regression analysis (Figure 6B), supporting that the cultivars containing green revolution alleles are vulnerable to temperature increase. Next, with another regression was performed on the YPP variations and the temperature variation based on the same dataset, showing an additional 4.8% and 7.5% yield penalty in cultivars containing *B1b* and *D1b* (Figure 6D), which are similar to the results from the regression on the YPP and average temperature (Figure 6B). Thus, the late sowing assay and regression analysis strongly suggest that the cultivars containing green revolution alleles are more sensitive to the temperature increase, will cause a more severe wheat yield penalty in the global warming scenarios.

## Discussion

Wheat as one of the most important crop, provides 20% food for human being ^27^. With the introduction of the dwarf genes by the Green Revolution, the yield of wheat was increased year by year ^28^. Up to now, about 70% of cultivars in the world carry one of the green revolution genes ^5^, which laid the foundation for wheat production supply for the increasing world population. Therefore, any agronomic traits and/or stress-resistance related to the green revolution alleles may influence the global wheat production and food supply security.

In this study, we clarified multitude agronomic traits controlled by the green alleles, which closely depend on the genomic background, and their phenotypes occur in filed with different probability. This result updates our understanding about crop phenotypes in field, they are not a fixed, but display a dynamic output depend on different genomic background. The understanding and global traits may provide helpful reference for precisely applying green revolution alleles in current and futural wheat breeding and production.

Global warming has resulted in large variations in wheat yield, with expected losses reaching up to 6% for each 1 °C rise in temperature ^29^ and warmer regions are likely to suffer more yield loss with increasing temperature than cooler regions ^30^. Compare with the preindustrial era, the global average surface temperature was about 1 °C higher ^31^, if worldwide carbon emissions continue at the current rate, global warming is likely to exceed 1.5 °C between 2030 and 2050, and even more than 3– 5 °C at the end of the 21st century ^32^. The situation is urgent to ensure the food supply with the global warming.

Generally, the higher temperature boosts the wheat development in booting, heading, anthesis, and maturity ^33,34^, and displaying a comprehensive alteration in wheat traits, consequent to a yield penalty (Figure 5 and Figure 6). For human, the yield is the most important target for sustainability. The yield of wheat is genetically determined by three individual components: spike number (the productive tiller number) per unit area, grain number per spike and kernel weight ^35^. Although the grain number per spike is significantly inhibited by the heat ^36,37^, the tiller number and kernel weight is more sensitive to the temperature increase (Figure 5 and 6), response to the major yield loss. Previous studies reported the productive tillers is significantly inhibited by the early high temperature by accelerate its development ^33,38^. Our data further showed that the tiller number reduction by heat contributes a major yield penalty, highlight the importance in maintaining the tiller number for securing wheat yield. In addition, the plant height is also highly sensitive to the temperature increase (Figure 5B and Figure 6A), further supporting that the biomass plays critical foundation for the grain yield and stress resistance ^18,39^. The higher biomass would be reconsidered in the wheat breeding program. Therefore, adjusting sowing time for bigger biomass and finding the heat-resistance QTL would be urgent mission for the global warming challenge ^39^ to maintain the wheat supply. Notably, the KNS is not sensitive to the temperature increase, implying the big spike with higher KNS may have the higher heat tolerance to avoid the yield penalty in the global warming scenarios, which may also be considered in current and futural wheat breeding program.

Our results showed the cultivars containing *Rht-B1b* and *Rht-D1b* have an increased KNS and kernel weight (Figure 2 and figure 6A), then still showed a significant lower YPP (Figure 6B), which would result from the significant reduced tillers number (Figure 6A). However, in wheat production, the cultivars containing green revolution alleles substantially increased the wheat yield per unit ^1,18^, implying a higher planting density were applied in cultivars containing green alleles. In this study, we also identified a significant decrease in flag leaf angle conferred by green revolution alleles, which would confer a more compact wheat plant structure, to allow a higher planting density and contribute to the significant yield increase per unit area. The higher planting density provides a plausible explanation for the yield increase of cultivars containing green revolution ^40-42^. However, these phenotypes controlled by green revolution alleles are sensitive to the temperature increase (Figure 5). The tiller number is the most heat sensitive traits for wheat yield components, showing a dramatic decrease with the temperature increase. To maintain the yield, the reduced tiller number require higher planting density for compensation. However, the flag leaf angle is also significantly increased with the temperature increase (Figure 5), which would prevent the higher planting density practices. Thus, the dilemma between tiller number and flag leaf angle in high temperature has to be resolved in futural wheat breeding. For the cultivars containing green alleles, especially containing *Rht-D1b*, the dilemma more prominent because both tiller number and flag leaf angle getting worse with the temperature increase, compared to the cultivars without green revolution alleles, which may further limit the production of cultivars with green alleles in the global warming scenarios.

Climate change may outpace current wheat breeding yield improvements ^30^, and more recently released winter wheat varieties were less able to resist high-temperature extremes than older varieties ^43^, we speculate that the increased frequency of the green revolution alleles in current wheat cultivars (Figure 4B) may partly response to the decreased heat resistance of current cultivars. To ensure the food security, we may consider more dwarf alleles for futural substitution, or improve the thermotolerant defect of green revolution alleles in time.

## Methods

### Plant materials and growing conditions

400 accessions of bread wheat were collected in this study, including modern cultivars and landraces from worldwide wheat accessions (Table S1). All materials were planted in seven environments (E1--E7) at Yangling (34°28′N, 108°07′E, altitude 517 m) and Chongzhou (30°63′N, 103°67′E, altitude 1300 m) of China in 2018-2021 (Table S11). Field experiments were arranged in a randomized complete block design with each variety planted in three replicates. Each accession was sown with 1 m in length and a row spacing of 20 cm, with 10 seeds per row. Field management was consistent with local practices with watering for wheat production.

Three F_2_ populations (Zhongluo08-3 × Ningmai12, Yannong22 × Qingfeng1 and MY72 × Changwei18) were planted at the Jize county of Hebei province (36°90′N, 114°80′E, altitude 42 m) in 2021-2022 growth season. Field management was consistent with local practices for wheat production.

### Agronomic traits measurements

The traits of plant structure and yield were measured at physiological maturity. Three plant individuals selected from each row were used to investigate plant height (PH), peduncle length (PL), total tiller number (TTN), productive tiller number (PTN), flag leaf angle (FLAN), spike length (SL), total spikelet number (TSN), fertile spikelet number (FSN) and kernel number per spike (KNS) according to established protocols ^8,44-46^. Days to heading was assessed as the interval between the date of seeding emergence and the date at which 50% of spikes per row emerged from the flag leaf, days to flowering was assessed as the interval between the date of seeding emergence and the date at which 50% of spikes per row have extruded at least one anther ^47^. Further, we calculated spikelet spacing (SS, SL/TSN) and yield per plant (YPP, PTN×KNS×TKW). In addition, thousand kernel weight (TKW), kernel length (KL) and kernel width (KW) were recorded using the rapid SC-G grain appearance quality image analysis system developed by Hangzhou WSeen Detection Technology Co, Ltd, China ^45^.

### DNA extraction and genotyping

Genomic DNA was extracted from wheat flag leaves of heading stage plants following the modified Cetyl Trimethylammonium Bromide (CTAB) method for genotyping ^48^. Kompetitive allele-specific PCR (KASP) markers were developed by the mutation site of *Rht-B1b* and *Rht-D1b* to identify all the accessions ^49^. All primers were listed in Supplementary Table S12. The KASP assay was performed as following: assays were set up as 5 μL reactions containing 2.2 μL template DNA dried to assay plate, 2.5 μL of V4.0 2x Kaspar mix (LGC Group, Teddington, UK), 2.444 μL ddH_2_O and 0.056 μL primer mix; and the PCR cycling included hotstart at 94 °C for 15 min, followed by 10 touchdown cycles (94 °C for 20 s; touchdown 61°C, -0.6 °C per cycle, 1 min), then followed by 35 cycles of amplification (94 °C for 20 s; 55 °C for 1 min).

### Genome-wide association study

The GWAS were performed on the SNPs derived population RNA-seq ^20^ with resample model inclusion probability ^50^. In details, 60% of each group were randomly selected without replacement and forward regression was performed, and repeated 100 times. The SNPs that were selected in the regression model in five or more subsamples were considered significant (RMIP ≥ 0.05) in this study. GWAS was implemented with the GAPIT version 3 in R (v3.6.1), using the FarmCPU ^51^ on each bootstrapped dataset. A Bonferroni-corrected threshold probability based on individual tests was calculated to correct for multiple comparisons using 1/N, where N is the effective SNP number calculated with the Genetic type 1 Error Calculator (v0.2) ^52^. The significant *P*-value thresholds for plant height and peduncle length were 8.58 × 10^−4^ in this population, but a more severe value *P* < 1×10^−4^ was selected to declare marker-trait associations (MTAs) in this study. To identify the traits that *Rht-B1b* or *Rht-D1b* are involved in regulating, those detected in at least two environments were considered to be stable identification.

### Statistical analysis

The best linear unbiased estimate (BLUE) values for phenotypic under seven environments were obtained using a mixed linear model by genotype as a fixed effect and replicates as random effects. Analysis of variance (ANOVA) was performed to test the homogeneity of genotype and phenotypic based on linear mixed model using the Python SciPy module. Differences between phenotypic of different genotypes were tested by *t-test* ^53^. If there is homogeneity of variance between the two sets of data, the default argument to the function *stats*.*ttest_ind* is used; otherwise, the parameter *equal_var* is set to *False*. All analyses were conducted using *lme4* (Bates et al., 2015) and *lsmeains* packages ^54^ in R3.6.2 ^55^.

### Permutation test

To reduce the interferes by co-selected SNPs and/or genes, the permutation test ^56^ was performed on each genotype pairs to evaluate the effect of different genomic background. The wheat population was divided into three groups by genotypes (*NO, B1b, D1b*), 50% of the accessions were randomly selected to test the significance between different groups. 1000 permutations were performed to examine the occurrence probability with significant differences between two groups.

### Multiple linear regression analysis of climate factors

Weather data were acquired from the China Meteorological Data Service Center (https://data.cma.cn). We collected 5 climate variables, including the maximum and minimum temperatures, sunlight hours, relative humidity and the solar radiation in seven environments. According to the heading and flowering time of each accession and 5 climate variables, the average temperature, the average sunlight hours, the average relative humidity and the average solar radiation at seeding stage, flowering stage and filling stage for each accession were collected according their developmental process (Table S7). Multiple regression analysis was used to examine the effect of meteorological factors on phenotypes ^25^. The equation was:

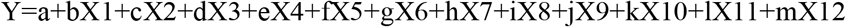

Where Y represents the phenotypic data. X1, X2, X3, X4, X5, X6, X7, X8, X9, X10, X11, X12, indicate the average temperature, the average sunlight hours, the average relative humidity and the average solar radiation at seeding stage, flowering stage and filling stage, respectively. b, c, d, e, f, g, h, i, j, k, l, m are the regression coefficients, and a is a constant. The Ordinary least square (*OLS*) was used for equation fitting, *P* < 0.05.

The *step* function of R3.6.2 was used for step-based regression, and the eliminated variables were selected according to Akaike Information Criterion (AIC). The meteorological factors with the smallest values from AIC were always removed until the equation was constructed successfully. The calculation formula of AIC: AIC=2k-2ln(L). L is the maximum likelihood function of the regression model. K is the number of parameters. The AIC of the model obtained by eliminating different variables were compared, the eliminated variables were selected in turn, and the final retained variables were ranked in importance from small to large according to AIC. The *vif* function of R3.6.2 was used to test whether the model has multicollinearity. The Variance Inflation Factor (VIF) was used to measure the severity of multicollinearity in multiple linear regression model, VIF<10, with no multicollinearity; 10 ≤ VIF<100, strong multicollinearity; VIF ≥ 100, severe multicollinearity. The error rate of multiple linear regression model was calculated by *sigma* function in R3.6.2.

### Multiple linear regression analysis of average temperature

Multiple regression analysis was used to examine the effect of average temperature at different stage on phenotypes. The equation was:

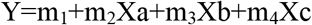

Where Y represents the phenotypic data. Xa, Xb, Xc represents the average temperature at seeding stage, flowering stage and filling stage, respectively. m_2_, m_3_, m_4_ are the regression coefficients, and m_1_ is a constant. The traits were predicted by 1 °C increase in the average temperature of all developmental stage.

The regression between YPP variations and the temperature variation. The variation of YPP and temperature can be obtained by subtracting BLUE in each environment. Based on the regression equation, the YPP was predicted by 1 °C increase in the average temperature of all developmental stage.

## Data availability

All data, models, or code generated or used during the study are available from the corresponding author by request. The phenotypes and SNP data are shown in https://iwheat.net/resource/. Meanwhile this data will keeps update every year, and we welcome more team adopt this population, phenotypes and meteorologic data, and may collect more information when we get more responses from users.

## Acknowledgements

This work was supported by “Integration of Two Chains” Key Research and Development Projects of Shaanxi Province “Wheat Seed Industry Innovation Project”; grants from the National Natural Science Foundation of China [31571756 and 31501380] and Key R&D Program of Yangling Seed Industry Innovation Center (Ylzy-xm-01). We thank Guoyue Chen and Zhien Pu for wheat field investigation in Sichuan Province, and thank Yufeng Zhang for wheat hybrid group field investigation in Hebei Province, and thank Bing Liu for the valuable comments to this study.

## Author contributions

S.X. designed the concept and experiments. Z.L., J.Z. and X.M. performed field investigation. J.C and X.Z performed the related germplasm identification. Z.L., Z.H., Z.Z., X.W. W.J., and S.X. analyzed the data and wrote the manuscript. All the authors were involved in the revision of the manuscript and approved the final manuscript.

## Additional information

**Supplemental Table 1**. The composition and phenotypes of 400 wheat accessions in seven environments.

**Supplemental Table 2**. GWAS of 13 wheat traits with RMIP (≥5).

**Supplemental Table 3**. The distributions of wheat accessions in different agroecological regions.

**Supplemental Table 4**. The duration days of all accessions at different stage.

**Supplemental Table 5**. The average temperature of different developmental stage in normal-sowing and late-sowing.

**Supplemental Table 6**. The alterations of wheat traits with late-sowing.

**Supplemental Table 7**. The meteorological data for all wheat accessions.

**Supplemental Table 8**. The heading and flowering time of all wheat accessions.

**Supplemental Table 9**. The multiple linear regression equations with multitude meteorological factors.

**Supplemental Table 10**. The multiple linear regression equations with average temperatures.

**Supplemental Table 11**. The filed investigation in seven environments.

**Supplemental Table 12**. The KASP primer sequence.

